# Human macrophages survive and adopt activated genotypes in living zebrafish

**DOI:** 10.1101/181685

**Authors:** Colin D Paul, Alexus Devine, Kevin Bishop, Qing Xu, William J Wulftange, Hannah Burr, Kathryn M Daly, Chaunte Lewis, Daniel S Green, Jack R Staunton, Swati Choksi, Zheng-Gang Liu, Raman Sood, Kandice Tanner

## Abstract

The inflammatory response, modulated both by tissue resident macrophages and recruited monocytes from peripheral blood, plays a critical role in human diseases such as cancer and neurodegenerative disorders. Here, we sought a model to interrogate human immune behavior *in vivo*. We determined that primary human monocytes and macrophages survive in zebrafish for up to two weeks. Flow cytometry revealed that human monocytes cultured at the physiological temperature of the zebrafish survive and differentiate comparable to cohorts cultured at human physiological temperature. Moreover, key genes that encode for proteins that play a role in tissue remodeling were also expressed. Human cells migrated within multiple tissues at speeds comparable to zebrafish macrophages. Analysis of gene expression of *in vivo* educated human macrophages confirmed expression of activated macrophage phenotypes. Here, human cells adopted phenotypes relevant to cancer progression, suggesting that we can define the real time immune modulation of human tumor cells during the establishment of a metastatic lesion in zebrafish.

---

Macrophages represent a mature population of terminally differentiated cells of myeloid-lineage found in all tissues(1, 2). They are often categorized by distinct functional properties, cell surface markers, and the cytokine profile of the microenvironment. Highly plastic, macrophages adopt diverse phenotypic and functional states to regulate tissue homeostasis, tissue patterning, branching morphogenesis, wound repair and immunity(2). They respond to environmental cues within tissues such as damaged cells, activated lymphocytes, or microbial products to differentiate into distinct functional phenotypes(3). However, macrophages may adopt functions that aid and promote disease due to environmental cues that arise as a result of abnormal physiological states such as obesity, fibrosis, brain neurodegenerative disorders and cancer(1, 4–7). In particular, one of the “hallmarks” of cancer and predictors of aggressive metastatic disease is the chronic presence of activated myeloid cells, such as tumor associated macrophages (TAMs), within primary tumors(8–10). Probing the role of the inflammatory response in the earliest stages of malignant transformation remains technically and ethically difficult in human subjects. Nevertheless, the broad importance of immune cell biology necessitates appropriate models to adequately study implications in human disease.

A number of efforts have been made to “humanize” animal model systems to study human homeostasis and disease *in vivo*, especially in the context of blood cancers(11–17). However, the role of hematopoietic stem cells (HSCs) and their derivative lineages, such as myeloid cells, have also been identified as regulators of tumor progression for solid cancers(4, 8). There are several classes of macrophages that may be found around tumors(2, 10, 18). One class of macrophages includes the classical “M1” macrophage phenotype, which is characterized by the production and secretion of elevated levels of pro-inflammatory cytokines, mediation of resistance to pathogens with strong microbicidal properties, high production of reactive nitrogen and oxygen intermediates, and evocation of Th1 responses. In contrast, M2 macrophages are characterized by their involvement in tissue remodeling, immune regulation, tumor promotion and efficient phagocytic activity(2, 10). A second class of macrophages that mediate metastasis, metastasis-associated macrophages (MAMs) is thought to drive therapeutic resistance and establish macrometastases at distant sites(19, 20). However, probing the role of the inflammatory response on the etiology of metastasis, namely when one or few cells have initiated the formation of a distant lesion in vivo, is challenging. Of all the animal models of oncoimmunology, mice remain the standard model, as one can interrogate human cells using immune-compromised strains and similarly examine syngeneic cancer cell lines in the appropriate immune competent strain(11, 16, 21). However, directly visualizing the colonization of multiple murine organs is expensive as it requires many mice to achieve sufficiently powered statistics. Simply put, a model system that recapitulates physiologically relevant aspects of metastatic disease in which cells can be observed at the single-cell level would profoundly benefit metastatic cancer research(22).

Zebrafish larvae are optically transparent at larval stages and conducive for non-invasive imaging *in vivo*. This, coupled with the fact that many processes involved in embryonic development, cell biology, and genetics are conserved across vertebrates, makes zebrafish an attractive model to study tumor-immune interactions during early stages of organ colonization(23, 24). In addition, organ architecture, composition and regulatory signaling networks for several organs, including the brain, are well conserved in embryonic zebrafish(25, 26). As early as the first few days after fertilization, key components of the Central Nervous System (CNS) such as neurons, astrocytes, and microglia, as well as specialized structures such as the blood-brain barrier and choroid plexus, have been identified in larval zebrafish. The innate immune system, comprised primarily of neutrophils, tissue specific macrophages and peripheral derived monocytes/macrophages, is also conserved at this early developmental stage (15, 25, 26). Hence, the embryonic zebrafish offers a system in which cell dynamics can be observed while sharing key cellular and structural characteristics with the mammalian organs. This model can be used to delineate tissue resident vs. recruited circulating monocytes in the context of inflammation and tumor education for either primary brain tumors or in the case of early metastatic colonization. However, a key feature missing in studies of immune response in zebrafish is a careful demonstration that human immune cells can be introduced in this system with conserved survival and cell response.

Here, we introduced human monocytes/macrophages into the zebrafish, both directly into circulation and in an organ-specific manner. We determined that both monocytes and macrophages (differentiated in culture) survive in innate immune competent zebrafish for up to two weeks post-injection despite the lower physiological temperature of the zebrafish (28.5°C). Similar results were obtained *in vitro*, where flow cytometry and RNAseq analysis revealed that human monocytes cultured at the physiological temperature of the zebrafish survive, differentiate, and show surface markers associated with mature macrophages in response to cytokines comparably to cohorts cultured at human physiological temperature (37°C). We also determined that human cells are transformed by zebrafish astrocytes by employing transgenic fish where fluorescently tagged astrocytes can be visualized *in vitro* and *in vivo*. Gene expression analysis of *in vivo* educated human macrophages revealed gene expression associated with activation. In summary, these results characterized the function of human immune cells in the *in vivo* environment and physiological temperature of *Danio rerio*. Thus, this model system allows for examination of the contribution of specific immune cells within specialized organ microenvironments.

## Results

Zebrafish are reared at the lower temperature of 28.5°C compared to 37°C, the physiological temperature of mammals(27). Zebrafish also have an innate and adaptive immune response beginning from two and nine days post fertilization (dpf), respectively. We thus asked if human immune cells can survive at the physiological temperature of zebrafish in the presence of the innate and immature adaptive immunity in an organ specific manner. Human primary monocytes that were differentiated into macrophages in culture were injected into the hindbrain of the zebrafish. Longitudinal imaging of the same fish one day and 7 days post injection (dpi; 3 dpf and 9 dpf, respectively) revealed that human macrophages survive for at least one week after injection and can be found dispersed in the brain **(Figure 1a-c)**. Similar examination of human macrophages injected into zebrafish where the host microglia are fluorescently tagged revealed survival and distribution within the brain **(Supplementary Figure 1a)**. We next examined the survival of an immortalized human monocyte cell line under the same conditions. Similar survival and morphologies were observed within the brain (**Supplemental Figure 1b)**. Examination of zebrafish two weeks post injection revealed that a few human macrophages can persist *in vivo* up to this time point **(Figure 1d, Supplemental Figure 1c)**. We next asked if the human cells migrated within the parenchyma when directly injected into the brain. Serial imaging revealed that human macrophages were widely dispersed within the zebrafish brain and were often found in close vicinity to blood vessels **(Figure 2**). As immune cells are involved in tissue remodeling and surveillance, we next asked if the introduced human cells show comparable motilities. We quantified the host immune cells movement by tracking neutrophils and macrophages in addition to introduced human monocytes within the brain at 3 dpf. We determined that neutrophils move with an average speed of 7 microns/minute, which was significantly greater than that of either the zebrafish brain resident macrophages (P<0.0001) and the introduced monocytes (P<0.0001). There was no significant difference between the slower monocytes/zebrafish macrophages (P=0.8263), where an average speed of 0.5–1 microns/minutes was observed **(Supplemental Figure 2)**.

**Figure 1.**
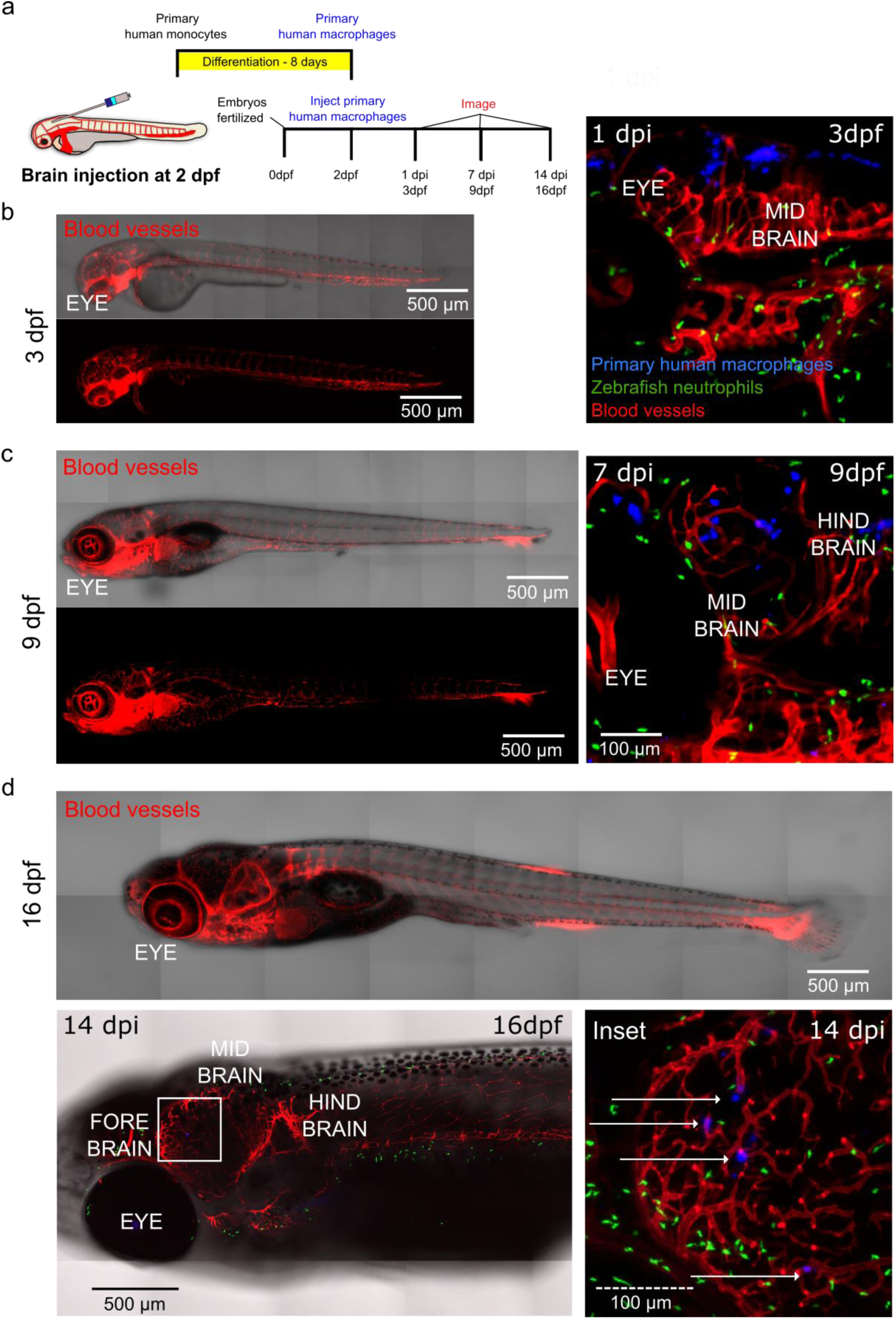
Human macrophages survive *in vivo* for up to two weeks postinjection following brain injection. (a) Schematic of experimental design: primary monocytes were differentiated into macrophages before injection into the zebrafish brain at age 2 days post fertilization (dpf) and imaged at 1, 7 and 14 days post injection (dpi). (b) Micrographs of representative whole larva at 3 dpf (left) and 3D projections showing distribution and survival of human primary macrophages (blue) injected into the hind brain of transgenic mpx:GFP (neutrophils-green)/flk:mCherry (vessels-red) zebrafish embryos at 1 dpi (3 dpf) (right). (c) Micrographs of representative whole larva at 9dpf (left) and 3D projections showing distribution and survival of human primary macrophages (blue) injected into the hind brain of transgenic mpx:GFP (neutrophils-green)/flk:mCherry (vessels-red) zebrafish embryos at 7 dpi (9 dpf) (right). (d) Micrographs show that cells can persist for up to 2 weeks after injection at 16 dpf. Top panel: representative zebrafish at 16 dpf. Left panel: micrograph shows tiled image of transgenic mpx:GFP (neutrophils-green)/flk:mCherry (vessels-red)16 dpf zebrafish, white square highlights region of interest in the zebrafish brain. Right panel: micrograph of the inset where the white arrows indicate human cells. Scales are indicated on each image.

**Figure 2.**
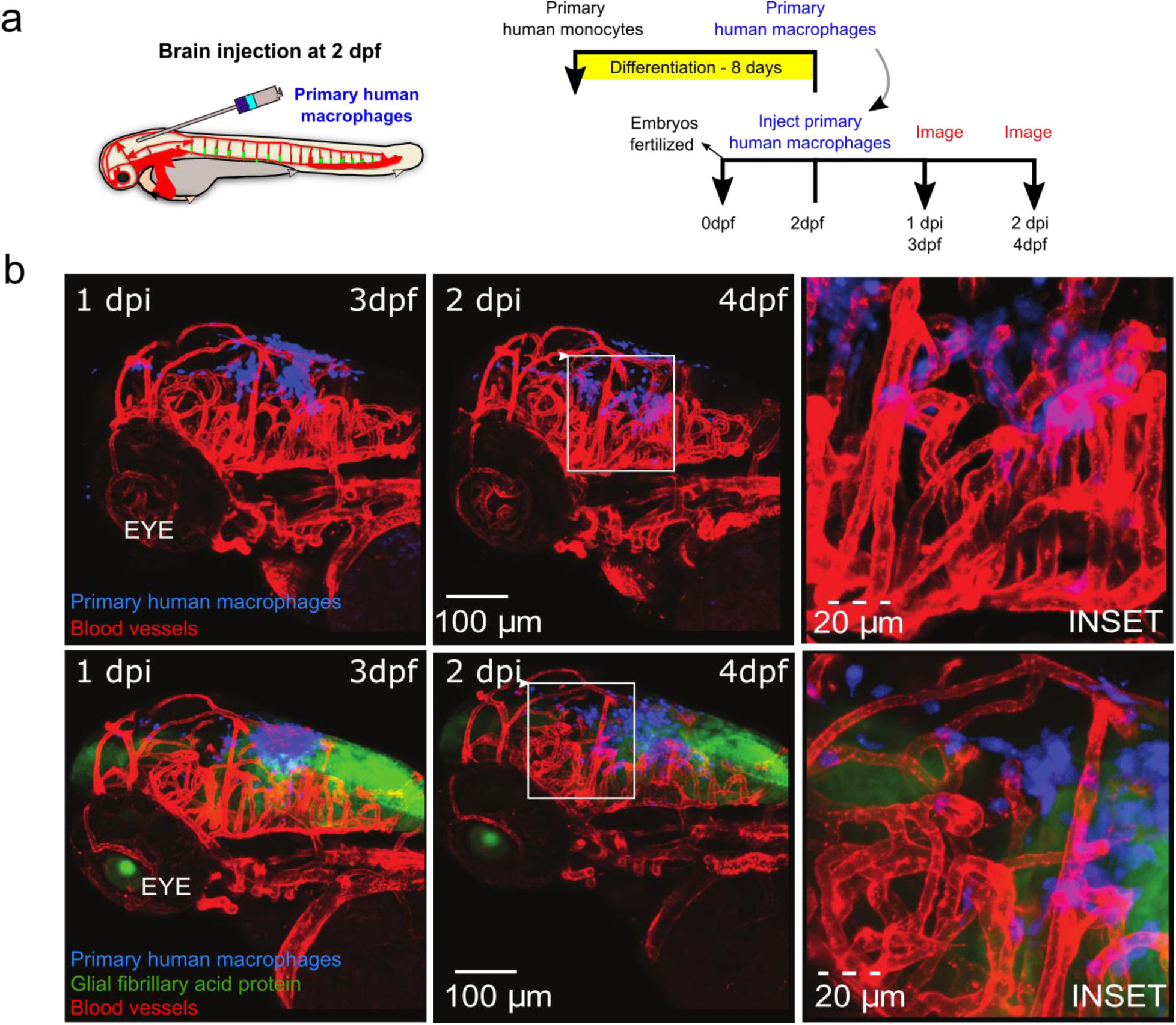
Human macrophages survive and interact with zebrafish stromal cells post-injection following brain injection. (a) Schematic of experimental design: primary monocytes were differentiated into macrophages before injection into the zebrafish brain at age 2 days post fertilization (dpf) and imaged at 1 and 2 days post injection (dpi). (b) 3D projections of whole head at 3 dpf (left) and 4 dpf (middle) showing distribution and survival of human primary macrophages (blue) injected into the hind brain of flk:mCherry (vessels-red) (top) and GFAP:GFP (astrocytes-green)/flk:mCherry (vessels-red) (bottom) zebrafish larvae. Boxes indicate positions of insets shown in right panels. Scales are indicated.

Immune cells are able to move from circulation to different tissues. Therefore, we asked if human cells would adopt similar phenotypes and survive in zebrafish upon access to the circulatory system. Human macrophages differentiated at either 28.5°C or 37°C and then injected directly to the zebrafish circulation were found throughout the entire zebrafish as early as ~3 hrs post injection and 1 day post injection **(Supplemental Movies 1–2, Figure 3a-c)**. They were observed to be motile and moved within intersegmental vessels and within different tissues **(Supplemental Movies 1–2)**. Imaging of zebrafish at one day and 7 days post injection (3 dpf and 9 dpf, respectively) revealed that human macrophages survive for at least one week after injection into the circulation and can be found dispersed in multiple tissues, including the gills, intersegmental vessels **(Figure 3d)**, and caudal tissue (data not shown). Interestingly, a subset of primary human macrophages injected to the zebrafish circulation at 2 dpf can disperse through the zebrafish and be found in the head and tail at 14 days post injection (**Figure 3e**).

**Figure 3.**
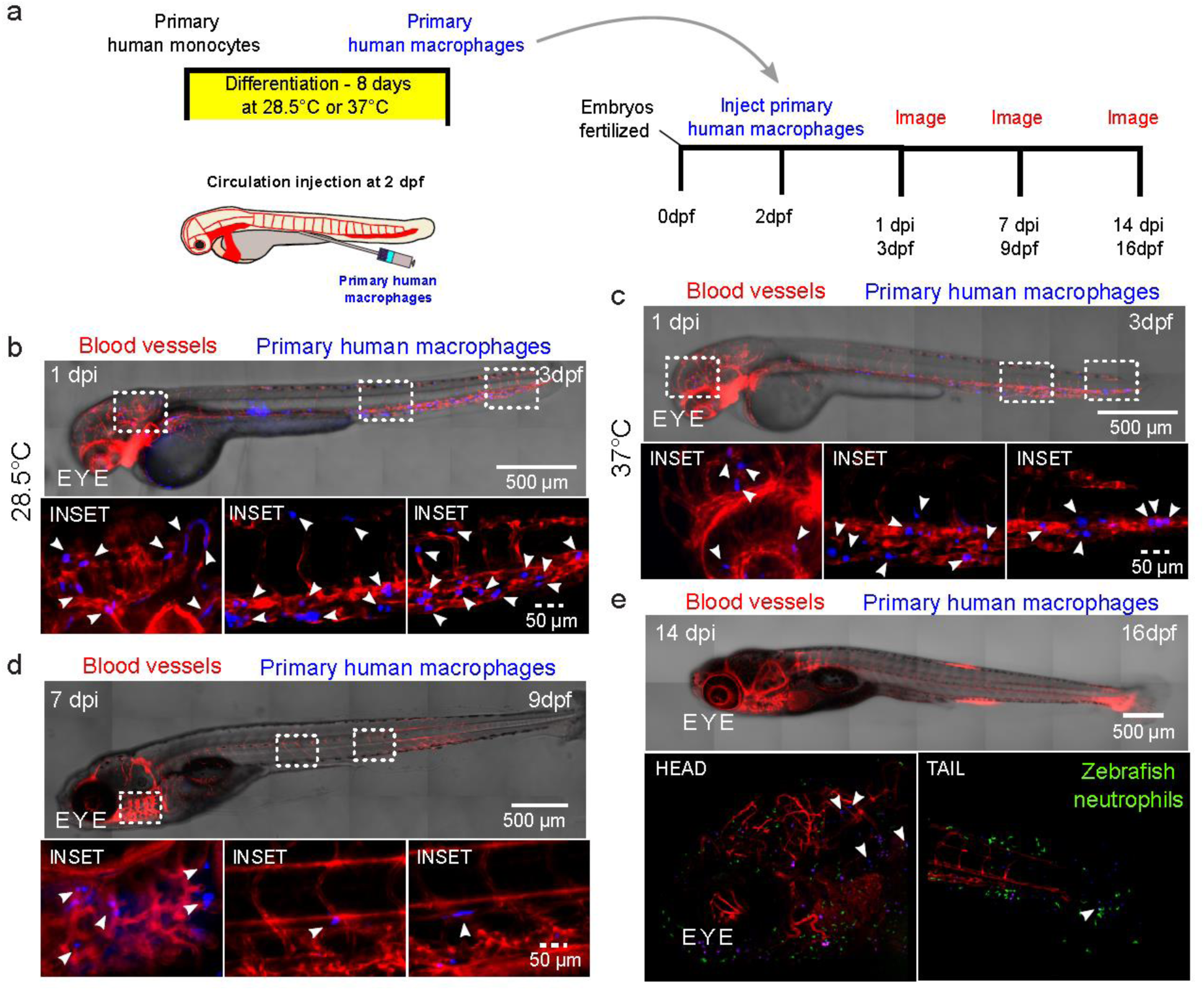
Human macrophages survive *in vivo* for up to two weeks postinjection following injection directly into the circulation of the zebrafish. (a) Schematic showing experimental set up, where human macrophages were directly injected into the circulation of 2 days post-fertilization (dpf) embryos, after being differentiated from primary human monocytes. Injected embryos were imaged at 1, 7 and/or 14 days post injection (dpi). (b-c) Micrographs of whole larva at 3 dpf (left) showing distribution and survival of human primary macrophages (blue) differentiated at either at physiological temperature of zebrafish (28.5°C) or at physiological temperature of humans (37°C) before injection into the caudal vein of transgenic fli:GFP (vessels-red) zebrafish embryos. Micrographs display macrophage distribution at 1 dpi. Bottom panels: Micrographs of 3D projections at higher magnification of insets highlighted in tiled images showing distribution and survival of human primary macrophages (blue). (d) Top panel: micrograph shows tiled image of the 9 dpf zebrafish, 7dpi, showing distribution and survival of human primary macrophages differentiated at human physiological temperature (37°C) (blue) injected into transgenic fli:GFP (vessels-red) zebrafish embryo. Bottom panel: Micrographs of 3D projections at higher magnification of insets highlighted in tiled image showing distribution and survival of human primary macrophages (blue). (e) Top panel: micrograph shows representative tiled image of a 16 dpf zebrafish. Bottom panel: Micrographs of 3D projections showing distribution and survival of human primary macrophages (blue) at 14 dpi following injection to transgenic mpx:gfp (neutrophils-green)/flk:mCherry (vessels-red) zebrafish embryos at 2 dpf. Arrows indicate macrophage positions.

Macrophages represent a class of terminally differentiated monocytes, and we reasoned that survival may differ as a function of differentiation status when cultured at the physiological temperature of zebrafish. Thus, we directly compared survival of human monocytes cultured at the two different temperatures. A subset of human monocytes were injected into the hind brain of the zebrafish, while others were cultured in vitro at either 28.5°C or 37°C. We imaged larva over the course of 3 days post injection at 24 hour intervals **(Figure 4a)**. Analysis of zebrafish imaged serially over the course of 1 day intervals revealed that 25–50% of cells survived in multiple transgenic lines after 3 days post injection (3 dpi) **(Figure 4b)**. There was no regional preference for survival as cells were observed throughout the fore, mid and hind brain of each animal **(Supplemental Figure 3a,b)**. Additionally, the zebrafish strain into which monocytes were injected was not a significant factor on monocyte survival (P=0.4917, **Supplementary Figure 3c**). We determined that a similar percentage of monocytes cultured at the zebrafish physiological temperature survived in vitro. Approximately 40% of monocytes from de-identified human donors survived at the zebrafish physiological temperatures and were viable up to 7 days of *in vitro* culture in only culture medium. In contrast, these same cells cultured at mammalian physiological temperatures showed rapid cell death and viability, where ~50% of cells underwent apoptosis after 2 days of *in vitro* culture in culture medium and only 10% of cells were present 7 days after plating **(Figure 4c)**.

**Figure 4.**
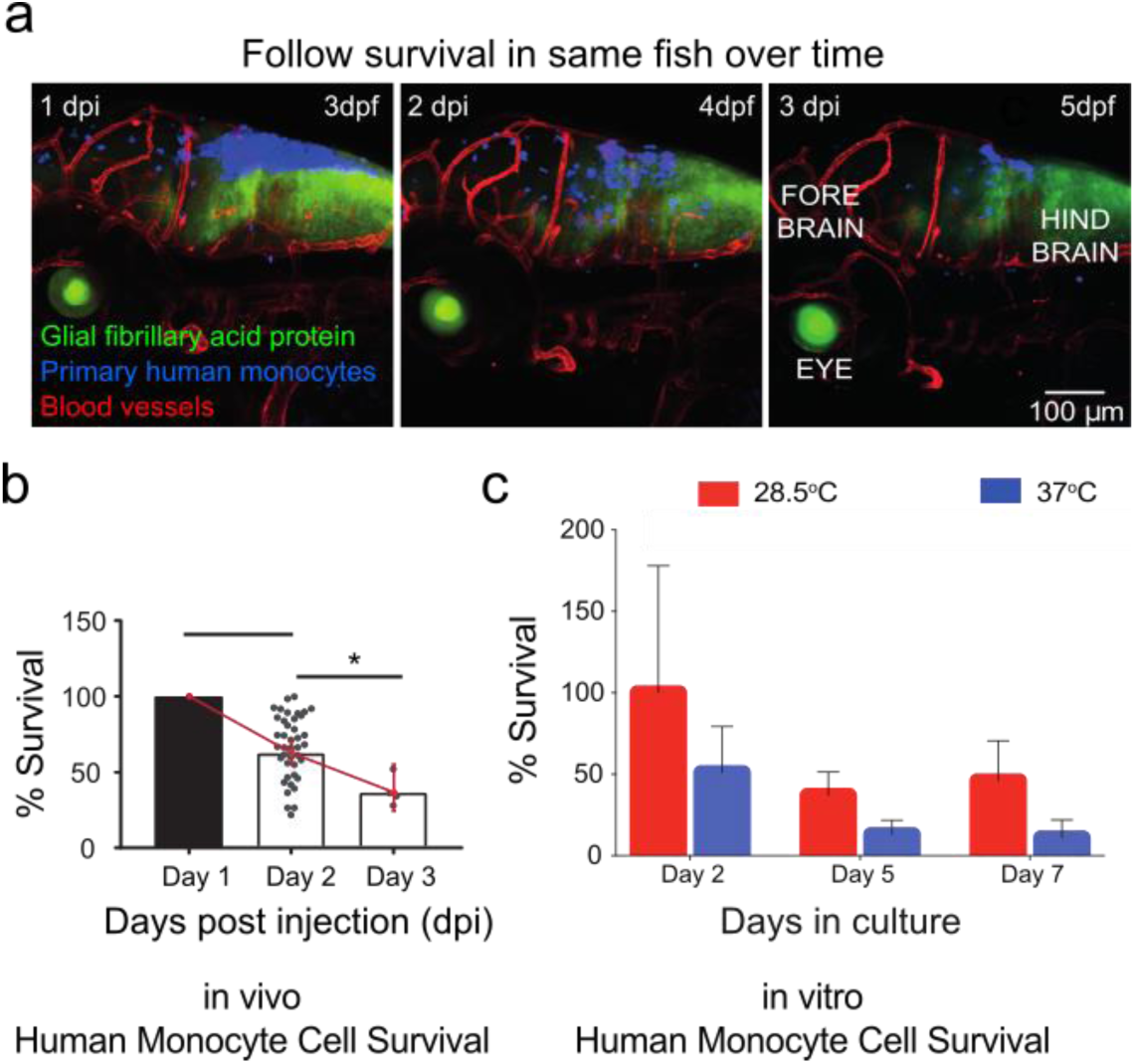
Primary human monocytes survive at physiological temperature of the zebrafish in vitro and in vivo. (a) Primary human monocytes (blue) were injected into the hind brain of transgenic GFAP:GFP (astrocytes-green)/flk:mCherry (vessels-red) zebrafish embryos at 2 dpf. Micrographs show representative images where zebrafish was imaged serially over the course of 3 days post-injection at interval of 24 hours. (b) Plot (mean±95% CI) of average in vivo survival calculated for primary human monocytes obtained from larvae where the numbers of cells that survived over the course of 3 days for each larvae were normalized to the initial numbers measured one day post injection. Statistical analysis where * indicates a p value of p<0.025 and *** indicates a p value of p<0.0005 by successive t tests with Bonferroni correction, with n=41 larvae for Day 1, n=41 larvae for Day 2, and n=4 larvae for Day 3. Scatter plots show values for individual zebrafish. Not all zebrafish survived to 5 days post injection. (c) Cells were either cultured at physiological temperature of humans (37°C) or at physiological temperature of zebrafish (28.5°C). Plot (mean±SD) of average survival calculated for primary human monocytes obtained from n=4 donors where the numbers of cells that survived over the course of 7 days were normalized to the original seeding density. Days in culture was a statistically significant factor in survival by two-way ANOVA with one degree of freedom (P<0.0001), as was the temperature of culture (P=0.0018).

We then investigated whether these human cells cultured at zebrafish physiological temperature are capable of differentiating into functional macrophages. Human monocytes differentiate into macrophages in response to *in vitro* addition of human macrophage colony stimulating factor (H-M-CSF)(28). We observed that monocytes cultured in the presence of H-M-CSF at the lower temperature were not adherent and remained spherical in suspension (**Figure 5a**). This spherical morphology was similar to that seen in control monocytes cultured without addition of H-M-CSF at both temperatures. In comparison, cells that were cultured at the higher temperature adopted the classical morphology of adherent elongated cells following treatment of H-M-CSF for five days **(Figure 5a)**. Immune cells respond to cytokines secreted by other cell types. To determine if monocytes/macrophages respond to zebrafish stroma, we co-cultured these immune cells with zebrafish astrocytes using transwell assays. Quantitation of cells that migrated in transwell assays revealed that a greater fraction of cells migrated in response to zebrafish astrocytes for each culture condition than under control conditions **(Figure 5b)**. We determined that monocytes cultured in the presence or absence of differentiation media showed an increased survival at the zebrafish physiological temperatures, where ~40% of cells without H-M-CSF, and ~70% of cells with H-M-CSF, were viable up to 5 days of *in vitro* culture. In contrast, these same cells cultured at mammalian physiological temperatures showed reduced viability when cultured with or without differentiation media **(Figure 5c)**. Macrophages derived from monocyte precursors adopt specific phenotypes depending on cues from the local tissue environment. We next analyzed cell surface markers using flow cytometry to monitor macrophage maturation. Cells that were cultured with H-M-CSF at both temperatures showed expression for the surface marker CD14 **(Figure 5d)**(29). This surface protein, CD14, is an indicator of macrophage maturation. Surface marker expression for differentiated human macrophages is observed after 5, 8 and 11 days of exposure for both physiological temperatures. However, when cells are cultured at the lower temperature, there is a temporal delay in the expression of CD14 compared to cells cultured at the higher temperature, and a subset of these cells remain CD14-negative **(Figure 5d)**. Monocytes that are subjected to M-CSF will differentiate into macrophage populations marked by expression of CD86, CD163 and CD206(30, 31). We thus asked if monocytes differentiated at the lower temperature will also show functional plasticity. Cells positive for CD14 expression were then assessed for markers, CD86, CD163 and CD206 at each temperature. Surface marker expression for differentiated human macrophages is observed after 5, 8 and 11 days of exposure for both physiological temperatures. However, when cells are cultured at the lower temperature, there is a temporal delay in the expression of CD86, CD163 and CD206, compared to cells cultured at the higher temperature **(Figure 5e)**. Functionally, monocytes differentiated into macrophages at the lower temperature were motile *in vivo* and showed similar motility to macrophages that were differentiated at the higher temperature **(Supplemental Movies 1 and 2)**.

**Figure 5.**
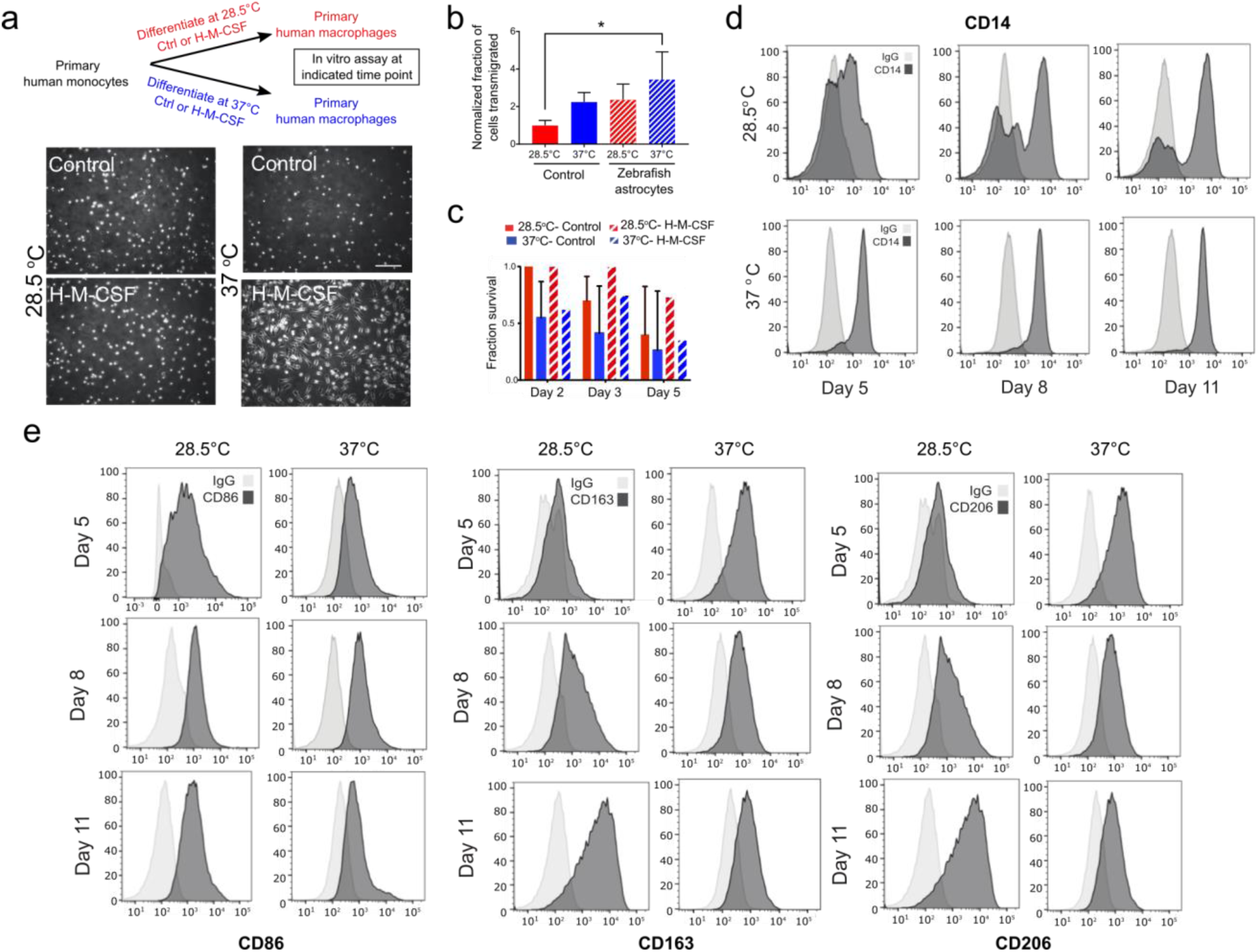
Flow cytometry determined that primary human monocytes can survive and differentiate into macrophages at physiological temperature of the zebrafish in vitro. (a) Schematic showing that human primary monocytes were cultured at physiological temperature of zebrafish (28.5°C) or at physiological temperature of humans (37°C) with or without differentiation by cytokine human macrophage colony stimulating factor (H-M-CSF) for 5, 8, and 11 days. Bottom panel shows micrographs show the morphology of cells in the presence or absence of cytokine at each temperature. (b) Graphs (mean±SD) showing fraction of migrated cells in response or absence of isolated zebrafish astrocytes in a transwell assay. *, P=0.0410 by Tukey’s multiple comparisons post-test, n=3 membranes/condition for astrocytes isolated on a single day. (c) Primary monocytes from 5 donors were either cultured at physiological temperature of humans (37°C) or at physiological temperature of zebrafish (28.5°C) with or without H-M-CSF. Plot (mean±SD) of average survival calculated for primary human monocytes/macrophages that survived over the course of 5 days were normalized to the original seeding density. Significantly more cells survived at 28.5°C vs. 37°C in the presence or absence of H-M-CSF by two-way ANOVA. (d) Human primary monocytes were cultured at physiological temperature of zebrafish (28.5°C, left panels) or at physiological temperature of humans (37°C, right panels) with cytokine human macrophage colony stimulating factor (H-M-CSF) for five, eight and 11 days. Expression of CD14 was assessed by flow cytometry. Light grey curve indicates control where shift in dark grey curve indicates expression. (e) Cells positive for CD14 expression were then assessed for markers, CD86, CD163 and CD206 by flow cytometry at each temperature and time point.

For the first three weeks of development the zebrafish innate immune system is the primary defense against invading pathogens. This is in contrast to mice, which have circulating T cells at 7 days post-partum(32, 33). Hence, we examined the innate immune response to the introduced human cells. Neutrophils are conserved in the axis of innate immunity in zebrafish(23). They form the first line of defense and produce a sustained inflammatory response following introduction of foreign entities. In the brain, the resident macrophages are the microglia(26). Innate zebrafish neutrophils and macrophages are present throughout 3 dpf zebrafish, including in the brain (**Figure 6a,b**). To determine if there was a differential zebrafish neutrophil or microglia response, differentiated human macrophages cultured at each temperature were injected into the hindbrain of 2 dpf zebrafish where either host neutrophils/ macrophages and vasculature were fluorescently labeled (**Figure 6c**). Serial imaging of the same zebrafish one day and 7 days post injection (3 dpf and 9 dpf, respectively) revealed that macrophages differentiated at each temperature survive for at least one week after injection and can be found dispersed in the zebrafish brain as observed in previous transgenic lines **(Figure 6d-f)**. We determined that similar numbers survived in either line (two-way ANOVA with one degree of freedom, P=0.4575 for interaction, P=0.3072 for temperature effect, P=0.4822 for zebrafish line effect; **Figure 6d)**, with survival rates comparable to those observed in previous experiments. Injected macrophages adopt elongated and spherical phenotypes and are seen along blood vessels and in the brain parenchyma **(Figure 6, Supplemental Figure 4a,b)**. No swarming of neutrophils was observed in the vicinity of the human monocytes at 1 day or 7 days post injection. Micrographs show that a few neutrophils are in close proximity to human macrophages, and neutrophil distributions are similar to those in uninjected embryos. We observed some co-localization between the host and human macrophages as determined by overlapping in the micrographs of the zebrafish imaged 1 day post injection. However, very little overlap is observed one week post injection.

**Figure 6.**
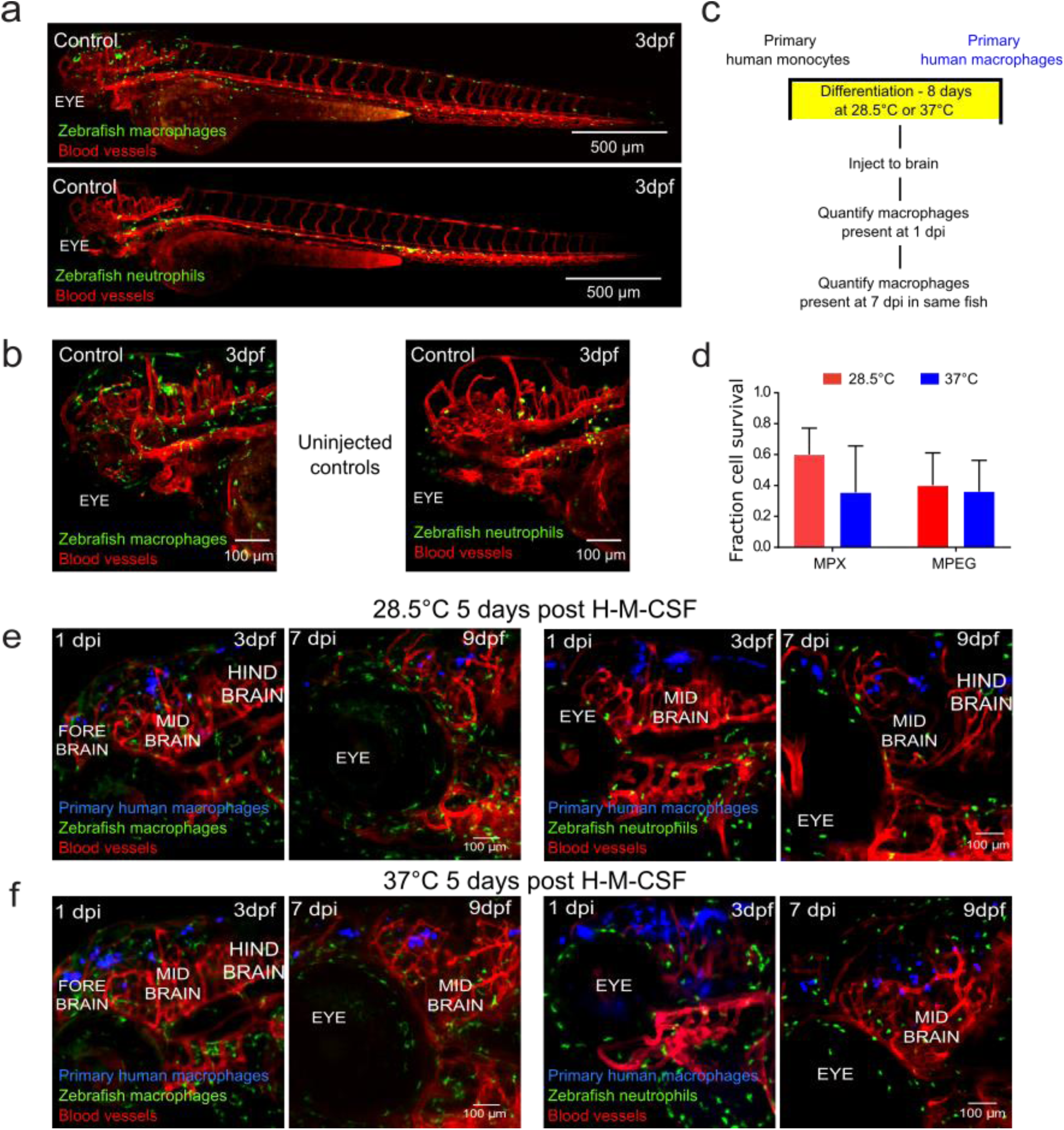
Primary human macrophages can survive at physiological temperature of the zebrafish in vivo and do not cause a sustained inflammatory response. (a) Representative light sheet micrographs of uninjected, control whole larva at 3 days post-fertilization (dpf). Both transgenic mpeg:GFP (macrophages-green)/flk:mCherry (vessels-red) (top) and mpx:GFP (neutrophils-green)/flk:mCherry (vessels-red) (bottom) are shown. (b) Representative lightsheet micrographs of head regions of uninjected, control embryos at 3 dpf. Both transgenic mpeg:GFP (macrophages-green)/flk:mCherry (vessels-red) (top) and mpx:GFP (neutrophils-green)/flk:mCherry (vessels-red) (bottom) are shown. (c) Schematic of experimental design: primary monocytes were differentiated into macrophages before injection into the zebrafish brain at age 2 dpf and imaged at 1 and 7 days post injection (dpi). (d) Plot (mean±SD) of average *in vivo* survival calculated for primary human macrophages differentiated at physiological temperatures of the zebrafish or human obtained from 3 larvae each, where the numbers of cells that survived over the course of 7 days were normalized to the initial numbers measured one day post injection. Differences in survival were not significant by two-way ANOVA with Tukey’s multiple comparisons post-test. The temperature of macrophage differentiation (P=0.3072), fish line injected (P=0.4822), and interaction between factors (P=0.4575) were not significant sources of variation by two-way ANOVA with one degree of freedom. Monocytes that had been cultured in the presence of human macrophage colony stimulating factor (H-M-CSF) for eight days (e) at physiological temperature of (28.5°C) and (f) at physiological temperature of humans (37°C). Primary human macrophages (blue) were injected into the hind brain of transgenic mpeg:GFP (macrophages-green)/flk:mCherry (vessels-red) or mpx:GFP (neutrophils-green)/flk:mCherry (vessels-red) zebrafish embryos at 2 dpf. Micrographs of 3D projections showing distribution and survival of human primary macrophages 1 dpi, when larvae are 3 dpf, and 7 dpi, when larvae are 9 dpf. The same fish is shown in the left and right panels.

In addition to tissue surveillance, macrophages remodel the tissue microenvironment. Thus, we asked if genes that encode for the matrisome were similarly expressed following in vitro differentiation at 28.5°C or 37°C. RNAseq revealed that 295/474 matrisome proteins were similarly expressed, with 179 others having statistically significantly different log10(normalized count) numbers. Notably, proteoglycan 2 was expressed at higher levels for macrophages differentiated at the physiological temperature of zebrafish. Moreover, examination of a cohort of genes associated with macrophage polarization revealed similar trends, with 71/164 genes similarly expressed and 93 having statistically significantly different log10(normalized count) numbers. Il-6 was expressed at higher levels for macrophages differentiated at the physiological temperature of zebrafish. Macrophages can polarize into different phenotypes in response to cues from the local tissue microenvironment(29). Thus, we compared gene expression of macrophages exposed to known cytokines in vitro at the two temperatures. We determined that similar expression of TNF-α, CD163 and VEGF for human macrophages cultured *in vitro* in the presence of recombinant EGF and TNF-α for 24 hours at both physiological temperatures. However, there was a greatly reduced expression of CCL18 for cells cultured at physiological temperature of zebrafish than at physiological temperature of humans **(Figure 7c)**. We then compared gene expression of human cells that were differentiated and then injected in the zebrafish brain for 24 hours prior to RNA purification.sfnalysis revealed similar expression of TNF-α and CD163 for human macrophages cultured in vitro at both physiological temperatures prior to injection. However, there was a reduced expression of VEGF, CCL17, CCL18, and CD24 for cells cultured at physiological temperature of zebrafish compared to physiological temperature of humans, though these differences were not significantly different. In contrast, IL 6 was expressed at higher levels in cells cultured at the physiological temperature of zebrafish than cells cultured at physiological temperature of humans, confirming the in vitro RNAseq result. **(Figure 7d).**

**Figure 7.**
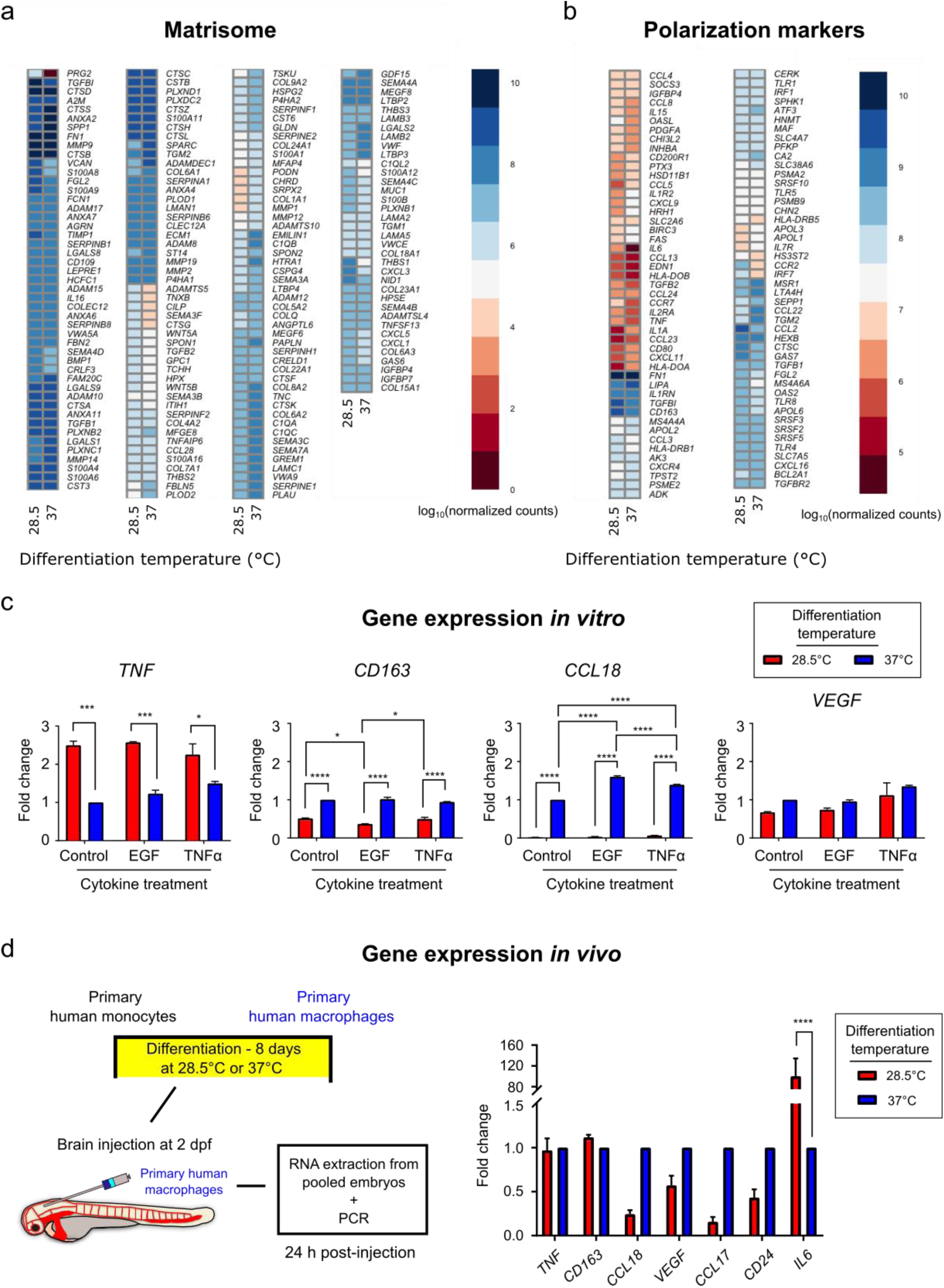
Primary human macrophages show gene expression of activated phenotypes in vitro and in vivo when cultured at the physiological temperature of the zebrafish. Human primary monocytes were cultured at physiological temperature of zebrafish (28.5°C) or at physiological temperature of humans (37°C) with human macrophage colony stimulating factor (H-M-CSF) for eight days prior to RNA isolation for RNAseq or further treatment with the cytokines EGF or TNFα prior to PCR. (a) Heatmap showing the log10 normalized counts of matrisome-associated genes. (b) Heatmap showing the log10 normalized counts for genes associated with macrophage polarization. In (a, b), heatmaps display genes for which p<0.05 and FDR<0.01 between temperatures. (c) Graphs show relative gene expression in specific markers for TNF–α, CD163, CCL18, and VEGF for human macrophages cultured *in vitro* in the presence or absence of recombinant EGF or TNF–α for 24 hours at physiological temperature of zebrafish (28.5°C) or at physiological temperature of humans (37°C) by real-time PCR. Plots are normalized to untreated macrophages differentiated at 37°C. (d) Human primary monocytes were cultured at physiological temperature of zebrafish (28.5°C) or at physiological temperature of humans (37°C) with human macrophage colony stimulating factor (H-M-CSF) for eight days prior to injection to the zebrafish brain. RNA was extracted from injected embryos 24 hours after injection. Graph shows *in vivo* relative gene expression in specific markers for TNF–α, CD163, CCL18, VEGF, CCL17, CD24, and IL-6 for human macrophages 24 hours after injection into the brain of the zebrafish. Plots are normalized to expression in macrophages differentiated at 37°C prior to injection and subsequent RNA isolation for each gene. For (c, d), *, p<0.05, ***, p<0.001, ****, p<0.0001 by two-way ANOVA with Sidak’s or Tukey’s multiple comparisons test.

## Discussion

Here, we introduce components of the human innate immune system to the zebrafish microenvironment by introducing human monocytes/macrophages. We first determined that human monocytes differentiate into functional macrophages at the physiological temperature of zebrafish. We showed that these human monocytes are able to survive for at least two-weeks post injection in vivo in immune competent zebrafish. Specifically, in transgenic zebrafish where fluorophores coupled to promoters of either myeloid-specific peroxidase *(mpx)*, a homologue of myeloperoxidase, or macrophage expressed gene 1 (*mpeg1*), allow specificity in distinguishing neutrophils from macrophages despite their common lineage. We used this distinction in lineage to analyze the survival and response of human immune cells introduced to embryonic zebrafish, demonstrating that human monocytes and macrophages injected into the zebrafish hindbrain or circulation survive and are functional for up to two weeks post-injection.

Introduction of human hematopoietic stem cells (HSCs) and hematopoietic cancers have been successfully performed in fetal sheep, immunocompromised mice, and zebrafish(10, 11, 16, 17, 24). In these animal models, the HSCs grafted successfully to the appropriate in vivo niche, bone marrow in mammals and kidney in the zebrafish, with varying lengths of survival in vivo. Moreover, the HSCs retained the ability to differentiate into multiple blood lineages, allowing the study of normal tissue homeostasis. This has greatly assisted in delineating what goes awry during transformation to cancers of the blood. Tissue grafting becomes more complicated in the case of cross species transplantation due to the mismatch in immune compatibility (11, 34). One solution involves sub-lethal irradiation of immune competent animals allowing for a temporal window whereby the donor components can be introduced (11, 34). However, there may be adverse and off-target effects due to the depletion of the recipient immune response. The zebrafish innate immune system comprised primarily of neutrophils and macrophages shows excellent conservation with mammals (15, 25). Here, we show that human monocytes/macrophages can survive in vivo in the larval zebrafish in the presence of innate immunity and immature adaptive immunity (**Figures 1–4, Supplemental Figures 1–3**). Interestingly, the introduced macrophages migrate through multiple tissues mimicking tissue surveillance, a key aspect of macrophage function **(Supplemental Movies 1 and 2)**. Furthermore, they co-exist seemingly unmolested by host neutrophils and macrophages **(Figure 6, Supplementary Figure 4)**. Thus, there is sufficient conservation in signaling proteins that regulate immune survival. This suggests that this model organism can provide a comparable study of metastases in the same manner as observed in mice between host tissues of zebrafish and human tumor cells.

In adult mammals, innate immune cells, monocytes and macrophages share a common myeloid progenitor in the bone marrow (2, 10, 11, 35, 36). These cells differ greatly in their lifespan, where peripheral blood derived monocytes (PBDM) survive just a few days before undergoing apoptosis. In contrast, differentiated monocytes such as macrophages may have a lifespan of months. While these immune subsets are conserved in zebrafish innate immunity, the major difference is that they exist at different physiological temperatures. This study showed that human monocytes also survive on the order of days at the physiological temperature of zebrafish and surprisingly, with a greater survival than a cohort cultured at the physiological temperature of human in vitro and in vivo **(Figure 4)**. Monocytes respond to different chemokines following a strictly regulated schedule (2, 10, 11, 35, 36). Following dissemination from the bone marrow, circulating monocytes are CCR2 negative. One example is that they become CCR2 positive roughly 1-2 days post release and successive tissue extravasation. Functional analysis revealed that human monocytes differentiate into M1 and M2 macrophages as dictated by high CD86, CD163 and CD206 positive expressions at both temperatures in a similar time frame. However, monocytes cultured at the lower temperature differentiate into mature macrophages with a temporal delay compared to monocytes culture at human temperatures. In addition, the cells displayed differences in morphology, where cells cultured at zebrafish physiological temperature remained in suspension and did not adhere, indicating that there are subtle differences due to the mismatch of culture temperature. One reason could be reduced metabolism at the lower temperature, or that the human cells follow the differentiation steps that zebrafish monocytes/macrophages adopt. Nevertheless, the functional status is achieved **(Figures 5 and 7)**.

Within a growing tumor, resident and recruited monocytes receive chemical cues from tumor-secreted chemokines and physical cues from the microenvironment such as hypoxia(37). Consequently, they become tumor-educated and adopt phenotypes that help drive tumor growth through promotion of neoangiogenesis, aberrant tissue remodeling, and development of immunosuppressive microenvironments(2, 4, 8). These tumor-associated macrophages (TAMs) are abundantly found within or adjacent to primary tumors in several types of human cancer. TAMs are regulated in part by colony stimulating factor (CSF)-1, mediated by the CSF-1 receptor (CSF1R), which drives macrophage maturation, tissue recruitment, and activation (38). Activated macrophages in turn release epidermal growth factor (EGF). At the primary tumor, this signaling pathway leads breast cancer cells and macrophages to form complexes that promote intravasation into the bloodstream (21, 39). There are several classes of macrophages that may be found around tumors. M1-type macrophages are thought to antitumorigenic and are characterized by high CD80, MHC-II, and PDL1 expression and low CD163 and CSF1R expression. In contrast, macrophages with high expression of CD68, CD163, CD206, and CSF1R are tumorigenic M2-type macrophages(20, 40) (18). Macrophages have been implicated in the clearance of circulating tumor cells during early metastatic dissemination (41, 42). Importantly, another class of macrophages, termed metastasis-associated macrophages (MAMs), may mediate metastasis, though the origins and recruitment of such macrophages are unclear(19, 20, 43). Here, we determined that in response to recombinant cytokines and brain-conditioning in vivo, human macrophages adopt gene expression that forms part of the identification of M1 and M2 phenotypes as measured by TNF-A, CD163 and VEG-F expression **(Figure 7)**. M2 macrophages have been classified into subdivisions, a, b and c(44, 45). The M2a subtype can also be defined as alternative activated macrophages, the M2b as type 2 macrophages, and the M2c as deactivated macrophages. The deactivated terminology refers to the in vitro ability of macrophages to adopt M2 activation following M1 activation, thus deactivating the M1-like gene transcription. Interestingly, CCL18 is differentially expressed as a function of culture conditions in vivo and in vitro **(Figure 7)**. CCL18 production is associated with the M2c subdivision for human monocytes and macrophages(45, 46). These data suggest that human monocytes differentiated at the lower temperature may not show the plasticity to switch between phenotypes compared to the macrophages differentiated at the human temperature. Interleukin-6 is a pleiotropic cytokine overexpressed in response to injury, inflammation, and infection(47). IL-6 is produced by many cells including osteoblasts, monocytes and macrophages(48). In our system, we observe that macrophages differentiated at lower temperatures express higher expression than macrophages differentiated at higher temperatures. Furthermore, this effect persists when cells are introduced in vivo. Interleukin-6 has a direct growth stimulatory effect on many tumor cells through the activation of several signaling pathways(48). Thus, in future studies, it may be more desirable to use human macrophages cultured at the physiological temperature of humans for co-injection of human tumor cells and macrophages.

Brain metastasis remains a challenge in clinical settings as patient survival is still measured in weeks or months (49). Emerging data indicate that immune-mediated signaling plays an important role in the establishment of brain metastasis (50). However, what is less understood is the role of tissue specific microenvironment of the immune cells at early stages of colonization restricted to lesions well below the current methods of detection. In a spontaneous mouse model of melanoma, not only were disseminated tumor cells detected before clinically detected primary tumors, it was determined that metastatic growth at distal organs was regulated by a tissue specific immune response (51). Recently, in studies of mouse models of brain malignancies and confirmed in patient data, that in addition to tissue resident macrophages microglia, an abundance of bone marrow derived monocytes (BDMs) can be found even in an immune privileged organ such as the brain (52). Furthermore, they uncovered that the resultant tumor associated populations were distinct based on the macrophage ontogeny within the brain. Here, we determined that human macrophages survive and become activated in the zebrafish brain environment. This model allows for examination of the contribution of immune signaling in a tissue specific manner. These findings open up the possibility of studying early metastatic events of human cancers in zebrafish. More importantly, it allows us to add the complexity of human tumor-human immune interactions whilst maintaining single cell resolution.

## Methods

### Animal studies

Animal studies were conducted under protocols approved by the National Cancer Institute, and the National Institutes of Health Animal Care and Use Committee. In our experiments, we employed several transgenic lines, i.e. Tg(mpeg1:GFP), Tg(gfap:GFP) and Tg(mpx:GFP), Tg(kdrl:GFP)la116 and Tg(kdrl:mCherry-CAAX)^y171^, kindly provided by Brant Weinstein (NICHD, Bethesda, MD), and the Casper strain, a kind gift from David Langenau (Harvard MGH)(53–55). Dual labeled vasculature and immune/astrocyte cells were generated by crossing Tg(kdrl:mCherry-CAAX)^y171^ and GFAP:GFP, mpeg:GFP or mpx:GFP. Progeny were screened using fluorescence microscopy to identify those that expressed fluorescent markers for vasculature and astrocytes, macrophages and neutrophils respectively. From this pool, founders were selected for continuation of the line. Zebrafish were maintained at 28.5°C on a 14-hour light/10-hour dark cycle according to standard procedures. Embryos obtained from natural spawning, raised at 28.5°C and maintained in egg water containing 0.6 g sea salt per liter of DI water were checked for normal development. At day 5 regular feeding commenced.

### Primary human monocytes-Maintenance and Macrophage differentiation protocol and functionality assays

Elutriated monocytes from 8 de-identified healthy human donors were acquired from the NIH, Department of Transfusion Medicine (DTM) (according to NIH protocol 99CC0168). Donors within the age group of 21–50 of either sex were selected for all experiments. Primary monocytes were cultured in DMEM high glucose, 10% fetal bovine serum and 1% penicillin/streptomycin. Cells were either cultured at physiological temperature of humans (37°C) or at physiological temperature of zebrafish (28.5°C). These cells were cultured at a concentration of 1.1×10^6^ cells /cm^2^ for minimum of five days to facilitate macrophage differentiation in the presence of recombinant human M-CSF (25 ng/ml). Media was replenished every 3 days. Samples were prepared for a minimum of three technical replicates for each donor sample. For comparison, a human monocyte cell line, U937, was cultured in suspension in RPMI media (ThermoFisher Scientific, Waltham, MA) supplemented with final volume of 10 % (v/v) Fetal Bovine Serum (FBS), Penicillin (100 U/mL) and Streptomycin (100 µg/mL). Cells were cultured at 37°C with 5% CO_2_ and relative humidity maintained at 95%. Media was refreshed every 2–3 days.

Primary monocytes that had been cultured in the presence of human M-CSF for eight days were cultured at each temperature (28.5 vs. 37°C). Cells cultured at each temperature were then cultured for an additional 24 hrs with a vehicle control and in the presence of either recombinant EGF at a concentration of 10 ng/ml or TNF-α at a concentration of 20 ng/ml. In addition, 2–5 nL of a sub set of the originally differentiated cells were then injected into 2 dpf fli:GFP embryos at a concentration of 1 × 10^7^ cells/mL for comparison. After 24 hrs, RNA was extracted for the *in vitro* educated macrophages. Similarly, RNA was extracted from pooled samples of 50 3 dpf embryos for each temperature (28.5 vs. 37°C) and age matched uninjected embryos.

### Quantification of in vitro monocyte survival at different physiological temperatures

For cell survival, human monocytes from 5 donors were seeded at a concentration of 1.1 × 10^6^ cells/cm^2^ and cultured at either 28.5°C or 37°C in base media. Human monocytes were collected and centrifuged at 1000 rpm for 5 minutes. Supernatant was removed and cells were resuspended in 1mL of sterile 1 × PBS. Cells were collected at 2, 3, 5 and 7 days after plating and analyzed for cell number and viability using Nexcelom Bioscience Cell Counter. Technical triplicates were collected for each donor sample for each temperature and day to obtain an average fraction survival per donor. Data were then displayed using GraphPad Prism 7.0a to graph cell number and viability as a function of days and temperature. Two-way ANOVA was used to determine statistical significance. Bright field and epi-fluorescent images were obtained on a Zeiss Axiovert 200 with an AxioCam MRm, using AxioVision LE 4.8.2 software. Images were acquired using 10× air objective with numerical aperture, 0.25, where each individual image had lateral dimensions of 1388 × 1040 square pixels corresponding to 0.448 × 0.335 mm^2^.

### Quantification of in vivo monocyte survival

Image processing was performed via ImageJ. Spectral crosstalk was removed by subtracting contributions of the red channel from the far-red detection channel. Images were then median (radius of 3 pixels) and Gaussian (sigma of 2) filtered to minimize background noise. Binary masks of monocytes were generated using Huang and Otsu thresholding methods on corrected images. The volume occupied by monocytes was then calculated using ImageJ’s built in ‘analyze particles’ function and the 3D shapes plugin. Number of monocytes was estimated by dividing the total volume of the binary monocyte mask by the expected average volume of a human monocyte (10 µm in diameter, volume of 104.7 µm^3^). Loss of monocytes was calculated for each larva by subtracting final number of monocytes from the initial number of monocytes (24 hours post injection) then normalizing by the initial number of monocytes. Percent survival was reported as 1 minus loss of monocytes multiplied by one hundred.

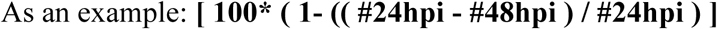

Data were then displayed using GraphPad Prism 7.0a to graph cell number and viability as a function of days. All error bars are 95% confidence intervals where D’Agostino and Pearson normality test was performed prior to either unpaired t-test or Kruskal-Wallis test to determine statistical analysis.

### Flow cytometry to determine cell surface expression to determine phenotype

Cells cultured in the absence and presence of human M-CSF were processed at minimum five days after plating for flow cytometry. Cells that remained in suspension were collected whereas adherent cells were detached with StemPro Accutase (ThermoFisher). For both conditions, the cell suspension was centrifuged and the supernatant removed as previously described. The cell pellets were re-suspended in ice-cold PBS with the following antibodies to assess surface expression for determination of macrophage polarization. Cells were incubated on ice with CD14-FITC (BD), CD86-PE (BD), and CD206-APC (BD), or with CD14-FITC, CD163-PE, and CD206-APC antibodies for 1 hour and cells were washed twice before processing for flow cytometry. Unstained cells and cells stained with single antibodies were used for gating cell size and setting instrument parameters. Samples were processed on a BD FACSCanto II (BD Biosciences) and analyzed with FlowJo software (Tree Star). PE- or APC-conjugated IgG stained cells were used to indicate background fluorescence and to set quadrants before calculating the percentage of positive cells.

### Fluorescence-activated cell sorting (FACS)

Embryos at 2 days post fertilization (dpf) with labeled vasculature and astrocytes cells were dechorionated in pronase solution (Roche Life Science #10165921001) as necessary (less than 5minutes). Embryos were then protease dissociated for FACS as previously described(56). Cell sorting was performed on a FACSAria Fusion instrument (BD Biosciences, San Jose, CA) using 85 µm nozzle at 45 psi pressure. Cells were gated on forward and side light scatter and separated based on GFP and/or mCherry-positive fluorescence signals defined with non-fluorescent negative controls.

### *In vitro* migration assays

Astrocytes isolated from FACS were centrifuged at 1000 rpm for 5 minutes, resuspended and counted. A 24-well plate was coated with 10 µg/ml laminin (Cytoskeleton Inc.) at room temperature for 1 h prior to triplicate wash with PBS. Fluorescence-activated cell sorted astrocytes were then added to the wells at 1 × 10^5^ cells per mL (1 ml/well in 24-well plate). Zebrafish astrocytes were maintained overnight in DMEM (Thermo Fisher) supplemented with 10% fetal bovine serum, 1% penicillin-streptomycin and 1% L-glut at 28.5°C. The following day, cells were prepared for transwell assay. Astrocytes were washed once with serum free RPMI medium. 700µl of serum-free media was placed on top of astrocytes. A sterile transwell membrane with 8 µm pores (Corning #3422) was inserted and 300µl of human macrophages were placed on top of membrane at 5 × 10^4^ cells per membrane. Transwells were incubated overnight at 33°C. Three membranes were run in parallel per condition (temperature and presence/absence of zebrafish astrocytes). Transwells were fixed in 4% paraformaldehyde for 10 minutes at room temperature. Membranes were washed twice in 1X PBS. They were permeabilized using 0.1% triton-x 100 for 5 minutes at room temperature, followed by a second wash step and then processed for immunofluorescence. Staining mixture consisted of Hoechst 33258 (10mg/mL, Thermo Fisher) was used at 1µg/mL in 1% BSA-PBS along with Phalloidin 565 (Sigma) used at 20µl/mL of Hoechst mixture. The membranes were imaged using Zeiss 780 LSM to acquire 5 × 5 tile scan images of each membrane, using a 10× air objective of 0.3 NA where each individual image comprised 2046 × 2046 square pixels corresponding to 1416 × 1416 square µm for a total Z distance of 100 µm with 5 µm step size. Three randomly selected 500 × 500µm fields of view were used for data analysis. We simultaneously excited our sample with the 405 nm from an argon ion laser with a power of<3 % (total power 30 mW) and 546 nm from a solid-state laser (power of< 10 %). A secondary dichroic mirror, SDM 560, was employed in the emission pathway to delineate the red (band-pass filters 560–575 nm) and blue channels (480–495 nm) at a gain of 400 on the amplifier. The laser power for the 543 nm setting was set at < 3 % of the maximum power and the gain on the detectors was set to 450. The Hoechst-stained cell nuclei were counted. The lower Z represents the bottom of the membrane and higher Z represents the top of the membrane. Nuclei counted as through were on the bottom or in pores. Nuclei counted on top were used in total cell count. Data were displayed using GraphPad Prism 7.0a to graph cell migration with given attractants. Two-way ANOVA with Tukey’s multiple comparisons post-test among all conditions was used to determine statistical significance.

### Zebrafish injections

For all experiments, at 24 hours post fertilization (hpf), embryos were transferred to egg water supplemented with phenylthiourea (PTU, Sigma P5272), suspended at 7.5% w/v in DMSO, at 1 part in 4500 to inhibit melanin formation for increased optical transparency. Embryos were then returned to the incubator at 28.5°C and checked for normal development. Zebrafish embryos at 2 days post fertilization (2 dpf) were anesthetized using 0.4% buffered tricaine. Human monocytes/macrophages were labeled with cell tracker (Green or Deep Red, Invitrogen) at a final concentration of 1 for 30 minutes. Cells were then centrifuged and the supernatant removed and pellet resuspended in PBS with a final cell concentration of 1 × 10^7^ cells/mL before injection. 2–5 nL of labeled cells were injected into the hindbrain of the embryo. Zebrafish were then reared at 28.5°C with 5% CO2 and relative humidity maintained at 95% for two days. At 5 days post fertilization, some zebrafish were then returned to system water and regular feeding at 28.5 °C for long-term survival studies. For injection to the circulation, zebrafish were anesthetized in 0.4% buffered tricaine and oriented in a lateral orientation on an agarose bed. Cells were injected directly in the circulation via the posterior cardinal vein using a pulled micropipette. Zebrafish were screened within 24 h of injection to check for successful introduction of cells to the circulatory system

### Intravital microscopy

For live cell imaging, embryos were anesthetized using 0.4% buffered tricaine, then embedded in a lateral orientation in 1% low melting point agarose (NuSieve GTG agarose, Lonza), and allowed to polymerize in with cover glass (no. 1.5 thickness) as previously described (57). Egg water supplemented with tricaine was added to the agarose hydrogel for the entire time of data acquisition. Zebrafish were imaged on Zeiss 710 or 780 laser scanning confocal microscopes. Initial overview experiments were taken at 20×, with 1 µm Z steps, as tile scans over the entire length and height of the zebrafish. Images in the head and tail of the zebrafish were acquired at 10× magnification every 10 min for 14 h. Z stacks were acquired using a tiled approach and a 10× air objective of 0.3 NA where each individual image comprised 2046 × 2046 square pixels corresponding to 1416 × 1416 square µm for a total Z distance of 276 µm. One-photon, confocal, 2-dimensional images of 512 × 512 pixels (lateral dimensions) were acquired with a 1.4 NA 40 × oil-immersion objective. We simultaneously excited our sample with the 405 nm, 488 nm lines from an argon ion laser with a power of < 3 % (total power 30 mW) and 546 nm from a solid-state laser (power of <10 %). A secondary dichroic mirror, SDM 560, was employed in the emission pathway to delineate the red (band-pass filters 560–575 nm) and green (band-pass filters 505–525 nm) and blue channels (480–495 nm) at a gain of 400 on the amplifier. The laser power for the 543 nm setting was set at<3 % of the maximum power and the gain on the detectors was set to 450.

### RNA isolation, RNAseq Analysis and Real-time PCR

Total RNA was extracted with Trizol according to the manufacturer’s guidelines (Invitrogen). Any remaining DNA was removed with the DNA-free kit (Ambion) and was re-purified with the RNAeasy kit (Qiagen). Illumina Stranded mRNA kit (Ref: 15056200) and the Illumina NeoPrep™ Library Prep System (Ref: 20003841) were used to prepare libraries from total RNA. Libraries were then sequenced using the Illumina NextSeq500 with 75-bp paired-end reads. Sequencing reads were trimmed for quality control using Trimmomatic (58) and quality was further verified with FASTQC (59). Trimmed reads were then aligned to human genome 19 (hg19) using STAR (60). Gene abundances were then determined using featureCounts (61) and normalized with TMM (62). Differential expression between samples was then determined using EdgeR (63, 64) and visualized with pheatmap (65). Genes associated with expression of proteins that comprise the matrisome and macrophage polarization were examined(66, 67). Genes plotted in Figure 7 were those from the matrisome and macrophage polarization profile for which p<0.05 and FDR<0.01. This cut-off was reached for 179/474 matrisome genes and 93/164 polarization genes.

Taqman real-time gene expression assays were run on an ABI StepOne Plus system according to manufacturer’s protocol (Applied Biosystems). Gene expression was normalized to that of GAPDH or β-actin. For analysis of gene expression in vitro, two biological replicates were used. Per biological replicate, cycle thresholds for each gene were normalized against a corresponding GAPDH replicate for each of two technical replicates, and technical replicates were averaged to determine a mean ΔcT and fold change compared to control macrophages differentiated at 37°C in the absence of EGF or TNFα. Results from the two biological replicates were then used to calculate the average fold changes and standard deviations. For analysis of gene expression from in vivo educated macrophages, cycle thresholds for each gene were normalized against a corresponding GAPDH threshold. Two technical replicates were then used to calculate mean fold changes and associated standard deviations. For each gene target, results were normalized to expression for in vivo educated macrophages differentiated at 37°C. All Taqman real-time primers used for gene-expression analysis were pre-designed and confirmed either from Integrated DNA Technologies: human TNFa (Hs.PT.45.14765639), VEGFA (Hs.PT.58.21234833), GAPDH (Hs.PT.39a.22214836), IL-6 (Hs.PT.58.39866843.g), CD24 (Hs.PT.58.39572682.g) or from Life Technologies: human CD163 (Hs 00174705), CCL17 (Hs01128674).

### Intravital cell tracking

Time-lapse microscopy images were exported to FIJI for analysis. To adjust for zebrafish growth during imaging, images of zebrafish injected with U937 cells were first registered using the Correct 3D Drift plugin, with the vasculature of the zebrafish used as a topographical reference. Innate immune cells in the head of 3 dpf Tg(mpx:GFP/flk:mcherry) or Tg(mpeg:GFP) zebrafish were imaged every minute. Immune cells were then tracked in 3D using the TrackMate plugin for FIJI. Images were segmented in the cell fluorescence channel in each frame using a Laplacian of Gaussian (14) detector with a 15 µm estimated particle diameter. An initial threshold of 1.0 was set, and the sub-pixel localization and median filter options in TrackMate were activated. Segmentation was then further refined by manual adjustment of the threshold to minimize the non-macrophage particles selected. Segmented objects were linked from frame to frame with a Linear Assignment Problem (LAP) tracker with 30 µm maximum frame-to-frame linking distance. Tracks were visually inspected for completeness and accuracy over the entire acquisition period and were manually edited to ensure that point-to-point tracks were generated for the entire time that a cell was in the field of view.

### Calculation of Cell Speed

Temporal and spatial information for each cell track was exported to MATLAB. For each cell, frame-to-frame speed was calculated by dividing the displacement of the cell by the time interval between frames. An average speed for that cell over the course of imaging was then calculated by averaging these frame-to-frame speeds. The ensemble of cell speeds for 1 zebrafish per condition (N=28 neutrophils, N=17 macrophages, N=13 monocytes) was compared in GraphPad Prism 7 using one-way ANOVA with Tukey’s multiple comparisons post-test following D’Agostino and Pearson normality test.

### Statistical Analysis

All statistical analysis was carried out in GraphPad Prism 7 (GraphPad Software, La Jolla, CA, USA). No statistical methods were used to predetermine sample size. Zebrafish were randomly selected from larger clutches prior to injection with human monocytes or macrophages. For analysis of *in vitro* monocyte survival, two-way ANOVA was used to determine statistical significance. Survival of monocytes *in vivo* for all zebrafish strains injected (monocyte survival data grouped for all zebrafish strains tested) was assessed by successive unpaired t tests, with significance levels adjusted by Bonferroni correction. For analysis of monocyte survival between gfap:gfp/flk:mCherry and mpeg:GFP zebrafish, an unpaired t test was used to assess monocyte survival following a D’Agostino and Pearson normality test. Similarly, survival of primary macrophages *in vivo* was assessed from 3 larvae per condition (temperature and zebrafish strain) by two-way ANOVA with one degree of freedom and Tukey’s multiple comparisons post-test. Macrophage transmigration through Transwell pores *in vitro* for three membranes per condition (temperature and presence of zebrafish astrocytes) was assessed by two-way ANOVA with one degree of freedom and Tukey’s multiple comparisons post-test among all conditions. To compare cell migration speeds *in vivo*, the ensemble collection of cell speeds were checked for normality using a D’Agostino and Pearson normality test, and average speeds between conditions were then assessed by Tukey’s multiple comparisons test. To assess gene expression in vitro by PCR, two-way ANOVA with Tukey’s multiple comparisons test was used across temperatures and cytokine treatments for each target. For PCR gene expression analysis following RNA isolation from macrophages in vivo, expression differences were assessed by two-way ANOVA with Sidak’s multiple comparisons test for each gene at 28.5°C and 37°C differentiation temperature. Statistically significant differences are indicated in figures with asterisks, where *, P<0.05, **, P<0.01, ***, P<0.001, and ****, P<0.0001.

### Data Availability

The authors declare that all data supporting the findings of this study are available within the paper and its Supplementary Information. All raw images and data generated in this work, including the representative images provided in the manuscript, are available from the corresponding author.

## Acknowledgements

This research was supported by the Intramural Research Program of the National Institutes of Health, the National Cancer Institute. We would like to thank Susan Garfield and Langston Lim, CCR Confocal Microscopy Core Facility, Laboratory of Cancer Biology and Genetics, NCI for use of the core microscopes. Cell sorting was performed by the NCI LGI Flow Cytometry Core. We would also like to thank the CCR Genomics Core, Center for Cancer Research, National Cancer Institute, National Institutes of Health. We would like to thank Ashley Williams for assistance in preliminary monocyte differentiation experiments. We would like to thank Zeiss microscopy for demonstration of a light sheet microscope used to generate Figure 6a,b using our samples.

## Author Contributions

K.T., Z.L, S.C, R.S., D.G. designed and discussed experiments, K.T., K.B., C.P., Q.A.D., C.L performed experiments, K.T., C.P., A.D., K.D., Q.X., C.L, J.S, H.B., W.W. performed data analysis. K.T., C.P., S.C. wrote the main manuscript text. K.T., C.P, Q.X., K.D., J.S., H.B., W.W. prepared figures. All authors reviewed the manuscript.

## Materials & Correspondence

Correspondence and material requests should be addressed to Kandice Tanner, Ph.D., 37 Convent Dr., Bethesda, MD 20852.

